# Dual functions of apigenin in suppressing *Phytophthora capsici* and shaping the pepper microbiome

**DOI:** 10.1101/2025.09.24.678249

**Authors:** Xiwen Liao, Weifeng Song, Yiman Wan, Jing Zhang, Panrong Ren, Yuan Chen

## Abstract

**Background:** Plant resistance to soil-borne pathogens is shaped by the interactions among host genetics, root exudates, and rhizosphere microbiomes. Flavonoids are widely recognized for their antimicrobial and signaling functions, yet their role in mediating metabolite-microbiome-pathogen interactions in pepper (*Capsicum frutescens*) remains poorly understood.

**Results:** Through integrated microbiome, transcriptome, and metabolome analyses, we compared resistant (CA53) and susceptible (CA476) pepper cultivars under challenge by *Phytophthora capsici*. Resistant plants maintained relatively stable transcriptional and metabolic profiles, whereas susceptible plants exhibited a pronounced suppression of the flavonoid biosynthesis pathway, with a marked decline in apigenin levels. Exogenous application of apigenin significantly enhanced pepper resistance by disrupting sporangial cell membrane integrity and thereby inhibiting zoospore release. In addition, apigenin functioned as a central hub metabolite, selectively enriching disease-suppressive rhizosphere microbes and reinforcing host protection.

**Conclusion:** Our findings uncover a dual role of apigenin in pepper resistance: directly inhibiting pathogen propagation and indirectly reinforcing the recruitment of protective microbiota. These insights highlight the ecological functions of root-derived metabolites in shaping plan-microbiome interactions and provide potential avenues for metabolite-informed strategies in sustainable crop protection.

## Introduction

Pepper (*Capsicum* spp.), a globally cultivated *Solanaceae* crop, holds significant economic importance due to its wide-ranging uses [1,2]. Despite the steady expansion of its cultivation, disease-related losses continue to threaten yield and quality. Among the major pathogens, *P. capsici* is a highly destructive soil-borne pathogen that causes root rot, foliar wilt, and fruit rot, posing a serious threat to pepper production worldwide [3]. Effective control of soil-borne pathogens remains challenging due to their persistent resting structures, such as oospores, and the presence of hidden initial infection sites within complex soil environments. Studies have shown that the rhizosphere microbiome in resistant plants is shaped by defense-related genes and contributes to disease suppression, nutrient uptake, and stress tolerance [4,5]. Growing evidence highlights the critical role of these beneficial microbes in strengthening plant immunity. As a result, manipulating the microbiome, especially those associated with resistant cultivars, has emerged as a sustainable and eco-friendly approach to disease control and improved crop productivity [6,7]. Host genotype plays a key role in shaping root architecture and influencing the composition of associated microbial communities, with numerous studies highlighting its strong impact on microbial recruitment. For example, resistant tomato plants selectively recruit two rhizobacteria, *Sphingomonas* sp. Cra20 and *Pseudomonas malodora* KT2440, which contribute to enhanced disease resistance in susceptible varieties [8].

Microbial recruitment is regulated by root exudates, which release diverse compounds that mediate biochemical signaling between plants and microbes [9,10]. Previous studies have shown that root exudates such as flavonoids and organic acids, play key roles in mediating plant–microbe interaction [11,12]. Root exudates from resistant and susceptible crop varieties differ in their effects on pathogens [13,14]. For example, methyl ferulate exuded by a resistant tobacco cultivar enhances resistance to *Phytophthora nicotianae* through a dual mechanism: directly inhibiting pathogen metabolism and indirectly recruiting beneficial microbes for synergistic suppression [15]. Interestingly, plants actively recruit specific rhizosphere microbes to cope with biotic and abiotic stresses, a process known as the “cry for help” strategy [16,17]. Through root exudates, plants emit distress signals that selectively attract beneficial microbes, reshaping the microbiome to enhance defense responses and stress tolerance. Recent studies have emphasized the vital role of root exudates in helping plants cope with both biotic and abiotic stresses [18–20].

Employing root exudate-mediated biological control provides a sustainable and efficient method for combating soil-borne pathogens. Flavonoids, as important components of root exudates, have been extensively studied for their role in the control of soil-borne diseases. Flavonoids can contribute to the control of oomycete diseases by interfering with the behavior of zoospores, particularly those of *Phytophthora* species. Specifically, certain flavonoids disrupt zoospore chemotaxis, which is the process by which zoospores recognize and locate host plants, thereby potentially reducing infection rates [21–23]. In addition to their role in defense against pathogens, flavonoids mediate selective cross-talk between plants and beneficial soil microbiota [24]. They may also influence microbial activity in the rhizosphere by modulating quorum sensing. For instance, Vandeputte et al. [25,26] demonstrated that several flavonoids, including apigenin, eriodictyol, kaempferol, luteolin, naringenin, naringin, and chalcone, affect the production of quorum-sensing-dependent factors in *Pseudomonas aeruginosa*.

Apigenin is a naturally occurring bioactive flavonoid synthesized in plants via the phenylpropanoid pathway. It is derived from phenylalanine or tyrosine through a series of enzymatic reactions catalyzed by PAL, C4H, 4CL, CHS, CHI, and FS, with intermediates such as cinnamic acid, p-coumaric acid, chalcone, and naringenin [27]. Liposome-encapsulated apigenin has been shown to significantly enhance antibacterial activity by inducing reactive oxygen species (ROS) generation in bacterial membrane lipids, increasing membrane permeability [28]. Recent studies have also demonstrated that apigenin and its derivatives can enrich *Pseudomonas* spp. in the rhizosphere of poplar, thereby promoting nitrogen utilization and lateral root development [29].

In this study, we hypothesize that resistant pepper varieties may induce or recruit beneficial microorganisms through specific root exudates following infection by *P. capsici*, thereby suppressing the pathogen and enhancing host resistance. To test this hypothesis, we analyzed the rhizosphere microbial communities associated with both resistant and susceptible pepper cultivars. By integrating metabolomic and transcriptomic analyses, we found that the flavonoid metabolite apigenin was significantly upregulated in the resistant cultivar compared to the susceptible one upon *P. capsici* infection. Further investigation revealed that apigenin directly inhibits *P. capsici* and contributes to the reshaping of the rhizosphere microbial community through root exudation. These findings shed light on the regulatory role of apigenin at the plant–microbe interaction interface.

## Results

### Screening of pepper cultivars for *P. capsici* resistance and exploration of rhizosphere microbiota and root-derived metabolites in regulating resistance

To assess variation in resistance, we screened 16 pepper cultivars against *P. capsici* using detached leaf and whole-plant inoculation assays, and identified eight resistant and eight susceptible cultivars (Fig. S1, Table S1). A resistant cultivar (CA53) and a susceptible cultivar (CA476) were selected for further study (Fig. 1A). Whole-plant inoculation showed an average disease index (DI) of 31% in CA53 and 73% in CA476 (Fig. 1B). Detached leaf assays revealed mean lesion areas of 86.64 mm² in CA53 and 345.78 mm² in CA476 (Fig. S1). These results confirm that CA53 is resistant, whereas CA476 is highly susceptible to *P. capsici*.

**Fig. 1.**
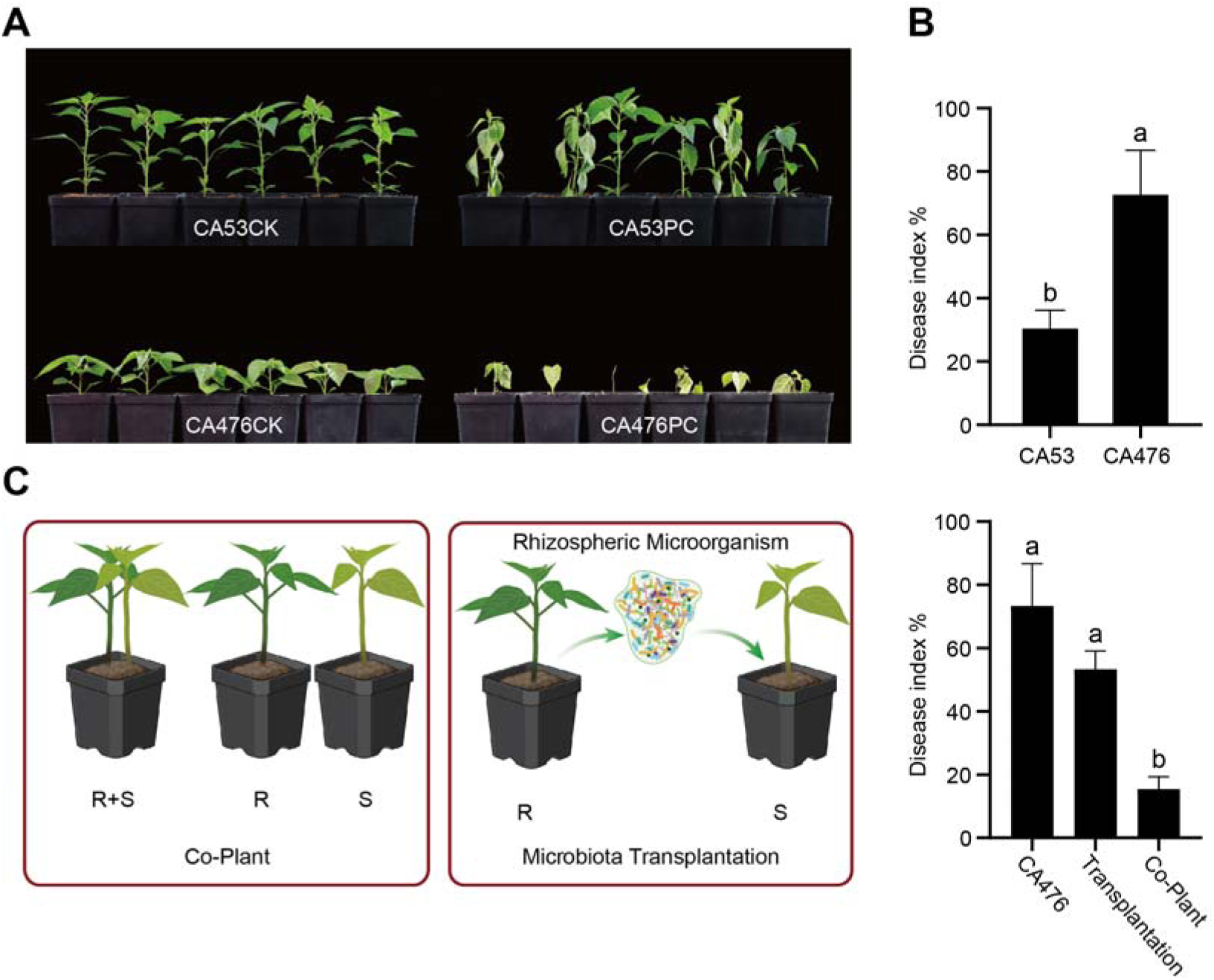
Pepper disease responses to *P. capsici* inoculation and microbiota-related treatments. (A) Phenotypic of disease-resistant and susceptible pepper cultivars after *P. capsici* inoculation. (B) Statistical analysis of disease index of the two varieties (disease-resistant vs. susceptible) in (A) after *P. capsici* inoculation. (C) Disease index of pepper under co-cultivation and microbiota transplantation treatments. Different lowercase letters (a, b) indicate significant differences among treatments (*P* < 0.05).

To investigate whether disease resistance in pepper cultivars is influenced by rhizosphere microbiota and root-derived metabolites, co-cultivation experiments were conducted using resistant and susceptible varieties. When grown together, the disease index of susceptible plants inoculated with *P. capsici* was significantly reduced, suggesting that resistant cultivars can enhance the defense of susceptible ones through rhizosphere-associated mechanisms. Microbiota transplantation further confirmed that transferring rhizosphere communities from resistant to susceptible cultivars suppressed disease development, though the effect was weaker than that observed in co-cultivation (Fig. 1C). This difference indicates that, beyond microbial contributions, root exudates of resistant cultivars may play a pivotal role in modulating disease resistance in susceptible plants.

Rhizosphere microbial and metabolomic divergence between resistant and susceptible pepper cultivars We first investigated the differences in rhizosphere microbial communities between resistant and susceptible cultivars. The richness index of rhizosphere microbial communities was significantly higher in resistant cultivars than in susceptible ones, suggesting that resistant cultivars harbor a broader spectrum of potentially beneficial taxa whose stability may contribute to disease suppression, whereas the Shannon and Simpson indices showed no significant differences between the two groups (Fig. 2A), indicating that inoculation with *P. capsici* did not markedly affect the richness or diversity of rhizosphere bacterial communities in either cultivar. Principal Coordinates Analysis (PCoA) based on Bray-Curtis distance revealed that the rhizosphere microbial communities of the disease-resistant cultivar (CA53) did not exhibit obvious separation before and after *P. capsici* inoculation, In contrast, the susceptible cultivar (CA476) exhibited a clear shift in community composition, as indicated by significantly separated confidence ellipses. These results suggest that inoculation had little impact on the rhizosphere microbiota of the resistant cultivar but markedly altered that of the susceptible cultivar.(Fig. 2B).

**Fig. 2.**
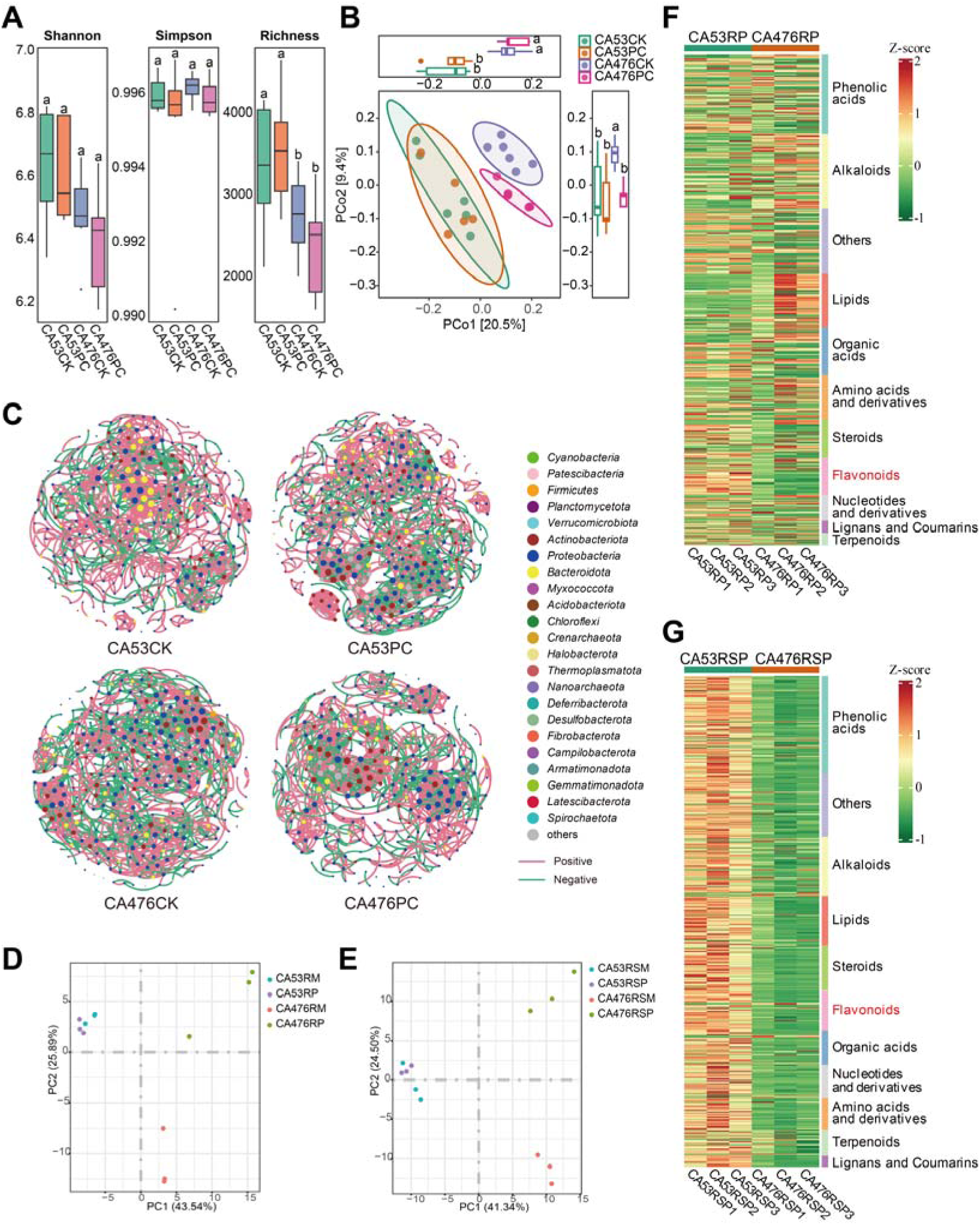
Rhizosphere microbiota and metabolites shifts in resistant and susceptible pepper cultivars after *P. capsici* inoculation. (A) Alpha-diversity indices (Shannon, Simpson, and Richness) of rhizosphere microbial communities. (B) Principal Coordinates Analysis (PCoA) of rhizosphere microbial community structures. (C) Co-occurrence networks of rhizosphere microbiota in four groups (CA53CK, CA53PC, CA476CK, CA476PC). CK: control treatment. PC: inoculation with *P. capsici* treatment. Nodes in different colors represent different bacterial phyla, and red lines indicate positive correlations while green lines indicate negative correlations. (D) Principal Component Analysis (PCA) of wide-targeted root metabolites. (E) PCA of wide-targeted root exudates. (F) Heatmap of root metabolites. (G) Heatmap of root exudates. RM represents root metabolites in the control group, RP represents root metabolites in the pathogen-inoculated group, RSM represents rhizosphere exudates in the control group, and RSP represents rhizosphere exudates in the pathogen-inculated group. Different lowercase letters (a, b) indicate significant differences among treatments (*P* < 0.05).

The rhizosphere microbial networks of resistant and susceptible cultivars also show great difference (Fig. 2C). The microbial network of resistant cultivars showed an increasing node number, edge connectivity, proportion of negative associations, average degree, and modularity after pathogen inoculation, whereas the same indices decreased in susceptible cultivars (Table S1). Thus, pathogen invasion enhanced the complexity of rhizosphere microbial networks in resistant cultivars but reduced it in susceptible ones. Previous studies have reported that greater microbial network complexity and a higher proportion of negative associations often contribute positively to community stability and resilience under external stresses [30–32]. This suggests that the reinforced and more complex network architecture in resistant cultivars may provide a buffering capacity against pathogen disturbance, thereby supporting sustained disease resistance.

To further elucidate the inhibitory mechanisms of resistant pepper cultivars to *P. capsici* infection, we performed a comparative analysis of the root exudates from CA53 (resistant, R) and CA476 (susceptible, S) inoculation with *P. capsici* inoculation using widely targeted metabolomics. A total of 1041 metabolites were identified in the samples of root exudates via UPLC-MS/MS (Fig. 2F, G and Table S2). Compound clustering analysis of root metabolit and root exudates revealed a clear distinction between the resistant line CA53 and the susceptible line CA476 after inoculated with *P. capsici.* (Fig. 2D, E).

Principal component analysis (PCA) of metabolomic data revealed that both root metabolites and root exudates of the resistant pepper line CA53 exhibited highly similar clustering patterns before and after pathogen inoculation (Fig. 2D). In contrast, the susceptible line CA476 displayed a pronounced separation in clustering following inoculation (Fig. 2E). These findings suggest that pathogen infection exerts only a minor impact on the metabolic state of the resistant line CA53, implying that its defense-associated metabolic networks may either preexist or rapidly reestablish stability. Conversely, the susceptible line CA476 undergoes substantial metabolic disruption after inoculation, which may compromise its resistance.

Heatmap analysis of root and root exudate metabolites revealed that, resistant pepper cultivars exhibited higher expression in multiple metabolic pathways after *P. capsici* infection, including those related to phenolic acids, amino acids and derivatives, flavonoid biosynthesis, and nucleotides, compared with susceptible cultivars. Notably, flavonoid biosynthesis was consistently enriched in both root metabolite and exudates of resistant lines, suggesting that flavonoids may play a central role in enhancing pathogen resistance (Fig. 2F, G).

### Transcriptome–metabolome integrated analysis reveals flavonoid-mediated resistance mechanisms

To validate this, transcriptome sequencing (RNA-seq) was performed on resistant line CA53 and susceptible line CA476 at two- and four-days post-inoculation. The results showed that flavonoid biosynthesis-related genes were broadly upregulated in line CA53 but downregulated in line CA476, further supporting the hypothesis that resistance is mediated through activation of the flavonoid biosynthetic pathway (Fig. S2 A-F). To elucidate differences in flavonoid metabolism between resistant (R) and susceptible (S) pepper lines in response to *P. capsici* infection, we integrated metabolomic and RNA-seq-based transcriptomic analyses. By combining flavonoid metabolite profiles with the expression of flavonoid biosynthetic genes, we constructed a relational network illustrating their interconnections. The results demonstrated significant differences in the expression of flavonoid biosynthesis–related genes between resistant (R) and susceptible (S) pepper lines after *P. capsici* inoculation. Specifically, seven *phenylalanine ammonia lyase* (PAL) and seven *4-coumarate-CoA ligase* (4CL) genes were markedly upregulated in R but downregulated in S lines, suggesting that enhanced phenylpropanoid metabolism may provide precursors for downstream flavonoid synthesis. Eight *chalcone synthase* (CHS) genes showed slight downregulation in R plants but stronger and widespread suppression in S plants. *Flavone synthase* (FNSL/FNSL) genes were transiently downregulated at 2 days but upregulated at 4 days in R lines, whereas they were consistently suppressed in S lines (Fig. 3A). Collectively, these expression patterns indicate that activation of the flavonoid biosynthetic pathway plays a pivotal role in enabling resistant peppers to withstand *P. capsici* infection.

**Fig. 3.**
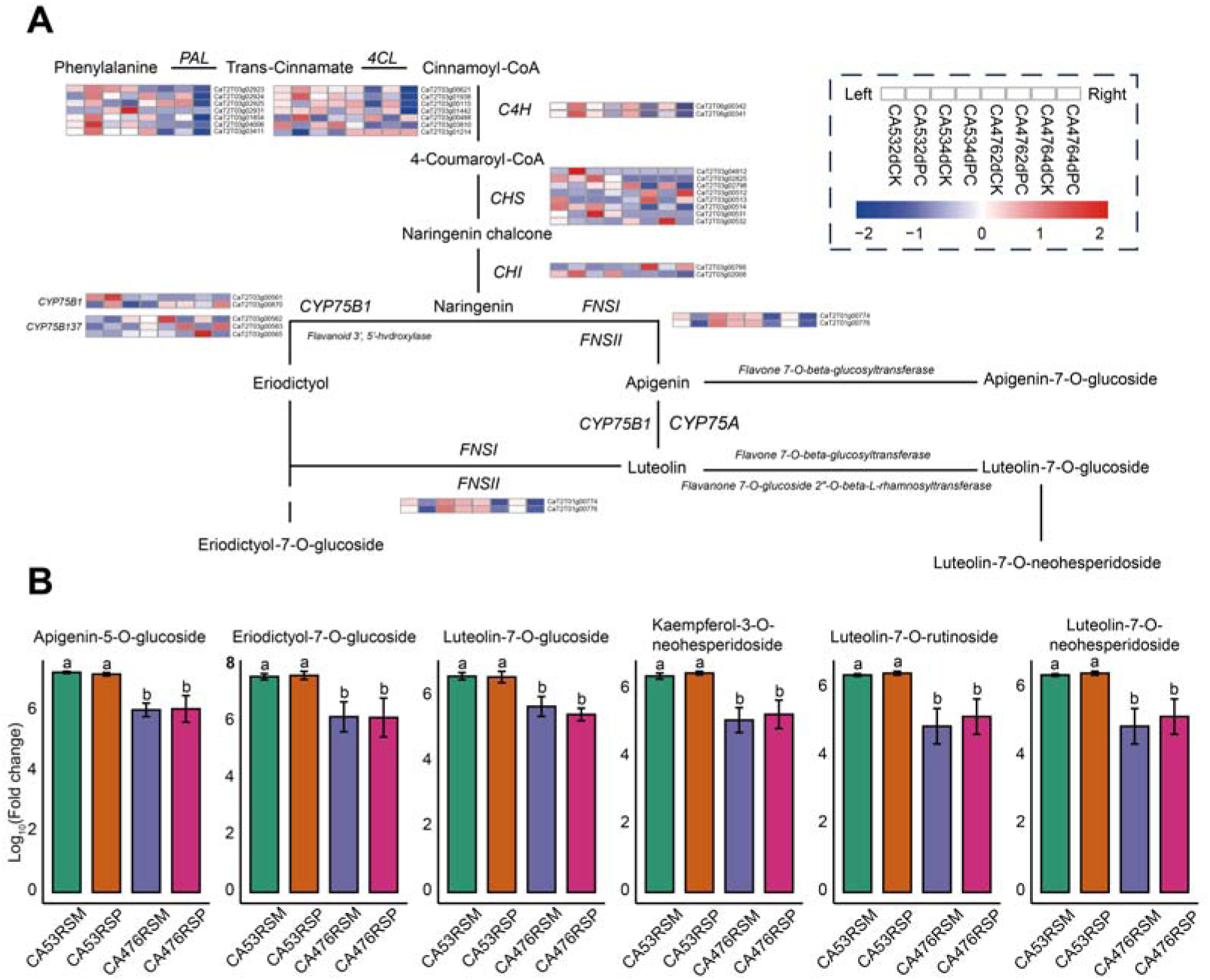
Differential expression of flavonoid biosynthesis genes and metabolite changes after *P. capsici* infection. (A) Expression pathway of 33 DEGs in response to *P. capsici* infection, which are involved in the flavonoid biosynthesis pathway (KO00941). The heatmap shows the expression patterns of the 33 DEGs in R and S accessions. White to red color represent upregulation; blue color represents downregulation. Heatmap expression values were standardized using Log_10_. Each sample had three biological replicates. (B) Box plots of the six flavonoid compounds, including Apigenin-5-O-glucoside, Eriodictyol-7-O-glucoside, Kaempferol-3-O-neohesperidoside, Luteolin-7-O-glucoside, Luteolin-7-O-neohesperidoside, and Luteolin-7-O-rutinoside, under different experimental groups. RSM represents rhizosphere exudates in the control group, and RSP represents rhizosphere exudates in the pathogen-inoculated group. Different lowercase letters (a, b) indicate significant differences among treatments (*P* < 0.05).

We compare flavonoid metabolomic profiling of resistant (R) and susceptible (S) cultivar in response to *P. capsici* infection, six markedly altered flavonoid compounds: apigenin-5-*O*-glucoside, eriodictyol-7-*O*-glucoside, kaempferol-3-*O*-neohesperidoside, luteolin-7-*O*-glucoside, luteolin-7-*O*-neohesperidoside, and luteolin-7-*O*-rutinoside were selected for further analysis (Fig. 3B). Subsequently, targeted metabolomic analysis was conducted to confirm the dynamics of these compounds and their precursors (apigenin and luteolin) and other structurally similar derivatives in root exudates. The results revealed that resistant peppers exhibited a significant accumulation of apigenin, luteolin, and their glycosylated derivatives (including apigenin-5-*O*-glucoside, luteolin-7-*O*-glucoside, and apigenin-7-glucoside), whereas no notable changes were observed in the susceptible line after *P. capsici* inoculation (Fig. S3). These findings indicate that the accumulation of flavonoids, particularly apigenin, luteolin and their glycosylated derivatives plays a critical role in the defense response of pepper against *P. capsici*.

### Flavonoid-mediated modulation of rhizosphere microbial interactions in resistant and susceptible peppers

We observed that flavonoid accumulation plays a pivotal role in the defense response of pepper against *P. capsici*. Previous studies have demonstrated that flavonoids secreted into the rhizosphere can influence microbial community structure and function, thereby promoting the recruitment of beneficial microbes and enhancing plant health and productivity [29,33]. To further elucidate how such flavonoid-mediated defenses coordinate with rhizosphere microbial dynamics to drive pepper resistance against *P. capsici*, we analyzed the composition of rhizosphere microbial biomarkers and their interaction networks following pathogen inoculation.

Distinct rhizosphere microbial assemblages were observed between the resistant and susceptible cultivars after pathogen inoculation, indicating adaptive microbial responses to invasion. To identify key taxa, LEfSe (Linear Discriminant Analysis Effect Size) was applied to both cultivars post-inoculation. The analysis revealed clear differences in biomarker enrichment: *Streptomyces*, *Pseudolabrys*, and *Brevundimonas* were enriched in the resistant cultivar CA53, whereas *Pseudomonas*, *Azospirillum*, and *Bordetella* dominated in the susceptible cultivar CA476. Notably, *Streptomyces* in CA53 and *Pseudomonas* in CA476 exhibited the highest LDA scores, suggesting their potential roles as critical determinants of resistance or susceptibility to *P. capsici* (Fig. 4A).

**Fig. 4.**
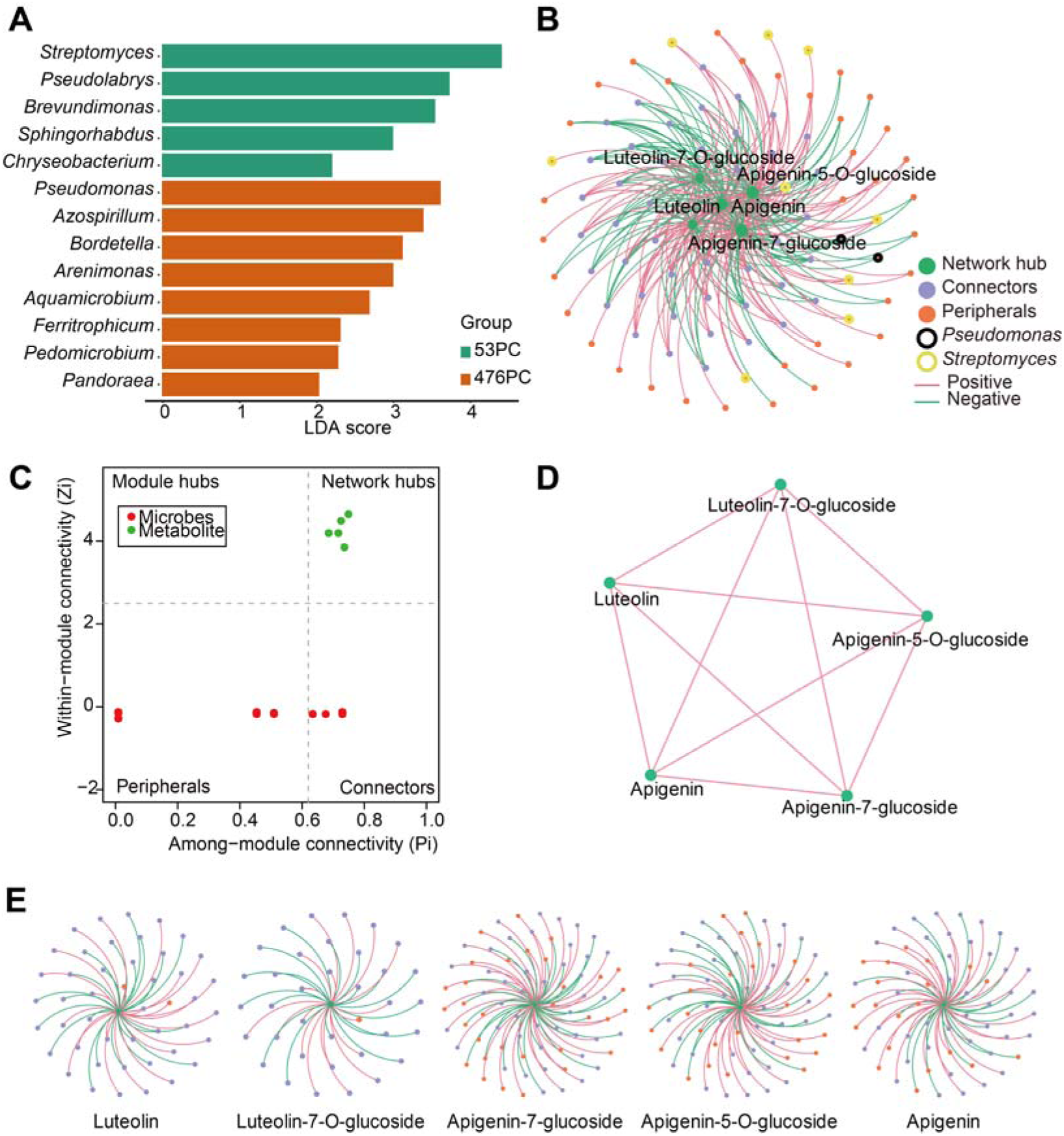
Network analysis of microbial communities and metabolites after *P. capsici* inoculation. (A) Linear Discriminant Analysis Effect Size (LEfSe) analysis showing differentially enriched microbial taxa between disease - resistant line CA53 (green) and susceptible line CA476 (orange) post - *P. capsici* inoculation, with LDA scores indicating the effect size of taxa differences. (B) Inter - Domain Ecological Network (IDEN) illustrating correlations between targeted metabolites (labeled nodes like luteolin, apigenin and their glycosides) and Amplicon Sequence Variants (ASVs, colored nodes). Red lines represent positive correlations, green lines negative correlations, and node colors distinguish network roles (e.g., green nodes as network hubs). (C) Scatter plot of within - module connectivity (Zi) versus among - module connectivity (Pi) classifying microbial nodes into network roles (peripherals, module hubs, connectors, network hubs).

To investigate the interactions between metabolites and microorganisms, we performed inter-domain ecological network (IDEN) analysis by integrating targeted detection data of five compounds (luteolin, luteolin-7-O-glucoside, apigenin, apigenin-7-O-glucoside, and apigenin-5-O-glucoside) with microbiome profiles. The analysis identified 89 ASVs associated with these metabolites, spanning 29 genera (Fig. 4B). All five compounds occupied hub positions in the network and exhibited significant positive correlations with one another (Fig. 4C, D). In addition, cultivar-specific microbial biomarkers also showed significant associations with these metabolites. In the resistant cultivar CA53, all ASVs belonging to *Streptomyces* were positively correlated with the metabolites, whereas in the susceptible cultivar CA476, ASVs affiliated with *Pseudomonas* were negatively correlated (Fig. 4B). This pattern implies that in CA53, *Streptomyces* may synergize with root-secreted metabolites (such as apigenin derivatives) to enhance resistance against pathogen invasion. By contrast, in CA476, *Pseudomonas* may act as a “helper microbe” for *P. capsici*; its abundance appears to be suppressed by metabolite accumulation, which could in turn weaken its facilitative role and thereby slow disease progression.

To further clarify the differential contributions of individual flavonoid types to microbial interactions, we conducted subnetwork analysis. Subnetwork analysis revealed that apigenin and its two glycoside forms were linked to markedly more ASVs compared with luteolin and its glycosylated derivative (Fig. 4E), suggesting that apigenin-related compounds may engage in more extensive potential interactions with the rhizosphere microbiota, highlighting their more prominent role in mediating the plant-microbe interactions underlying pepper resistance to *P. capsici*.

### Apigenin enhances pepper resistance by inhibiting zoospore release of *P. capsici*

Subsequently, apigenin, luteolin and apigenin-7-glucoside were used to validation assays to evaluate its inhibitory effects on *P. capsici* zoospore formation. The results demonstrated that all three tested flavonoids significantly reduced zoospore production, with apigenin showing the most pronounced effect, leading to a 90% reduction in zoospore numbers compared with the control (Fig. S4A, B and Supplementary video1-4). These findings demonstrate that apigenin potently suppresses the release of zoospores, thereby neutralizing the primary inoculum responsible for infection. We next examined whether apigenin affect the infection of *P. capsici* in pepper plants. In the detached leaf assay of the susceptible line CA476, application of three flavonoids (apigenin, luteolin, and apigenin-7-glucoside) followed by inoculation with *P. capsici* zoospores (10L spores/mL) showed that only apigenin significantly reduced lesion size, whereas the other compounds did not exhibit noticeable effects (Fig. S4C, D). Consistent with the zoospore release assay, apigenin stood out among the tested flavonoids, and was further evaluated *in vivo*. Whole-plant inoculation assays revealed that treatment of CA476 with apigenin (1 or 2.5 μg/mL) prior to root inoculation with zoospores (10L/mL) markedly reduced the disease index (DI) at 7 dpi and strongly inhibited the progression of root necrosis (Fig. 5A, B). Together, these results demonstrate that apigenin effectively suppresses the pathogenic process of *P. capsici* and significantly enhances pepper resistance to phytophthora blight.

**Fig. 5.**
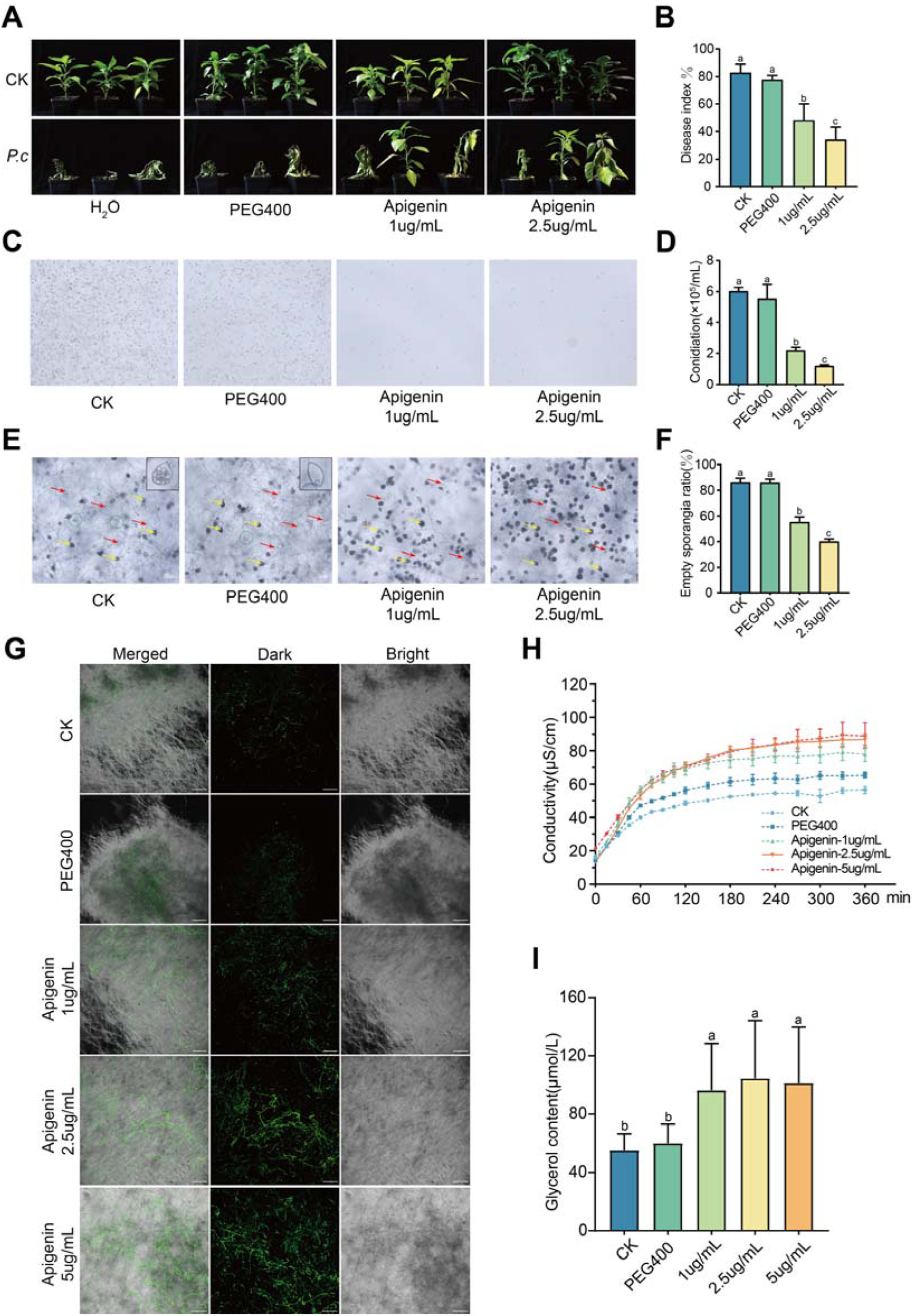
Apigenin improves resistance in susceptible pepper cultivars against *P. capsici* by inhibiting zoospore release and altering cell membrane permeability. (A) The effect of different apigenin concentrations on the *P. capsici* resistance phenotype of susceptible pepper (CA476). (B) Disease index of the peppers with apigenin before inoculation with *P. capsici*. The error bars indicate the standard deviations of three replicates. (C) The impact of different concentrations of apigenin on zoospore release. (D) Statistical of the release of zoospores after apigenin treatment. (E) Following treatment with apigenin, the phenotypic characteristics of sporangia in *P. capsici* were analyzed. Empty sporangia are annotated with red arrows, non-released sporangia with yellow arrows, and structures undergoing sporangiospore release are denoted by green circles. Bar=100μm (F) Quantitative analysis of empty sporangia(G) SYTOX^TM^ Green stain was employed to assess of *P. capsici* mycelia following apigenin treatment. Bar=100μm. (H) Measurement of conductivity of *P. capsici* following apigenin treatment.(I) Determination of glycerol content in *P. capsici* after apigenin treatment. Different lowercase letters (a, b) indicate significant differences among treatments (*P* < 0.05).

To investigate the mechanism by which apigenin inhibits zoospore release, apigenin was added at 1 μg/mL and 2.5 μg/mL during zoospore induction, and zoospore release was dynamically observed using light microscopy (Fig. 5C, D and Supplementary video5-8). Quantification based on empty sporangia (marked with red arrows) revealed that apigenin treatment significantly reduced the proportion of empty sporangia, accompanied by a concurrent decrease in both un-released (yellow arrows) and actively releasing sporangia (green circles), with the inhibitory effect increasing at higher concentrations (Fig. 5E, F).

During the zoospore induction stage, we assessed cell membrane integrity and permeability following apigenin treatment. Compared with the control, hyphae treated with increasing concentrations of apigenin (1, 2.5 and 5 μg/mL) exhibited strong green fluorescence, with fluorescence intensity significantly enhanced, indicating compromised membrane integrity (Fig. 5G). Consistently, conductivity assays showed that membrane permeability of *P. capsici* hyphae was markedly increased after apigenin treatment (Fig. 5H). In addition, apigenin treatment effectively activated osmotic regulation by enhanced glycerol biosynthesis and intracellular glycerol accumulation (Fig. 5I). Together, these findings suggest that apigenin may inhibit zoospore release by targeting membrane function and disrupting membrane permeability and osmotic homeostasis in *P. capsici*.

### Transcriptomic analysis reveals apigenin targets membrane permeability and ABC transporters

To further investigate the of apigenin suppresses zoospore release in *P. capsici*, we performed transcriptome sequencing and analysis of hyphae at the zoospore release stage following apigenin treatment. A total of 1,535 differentially expressed genes (DEGs) were identified, including 276 upregulated and 1,259 downregulated genes (Fig. 6A). KEGG pathway enrichment analysis revealed that these DEGs were significantly enriched in the ABC transporter pathway (Fig. 6B). ABC (ATP-binding cassette) transporters are key transmembrane components involved in the transport of diverse substrates [34]. Among the 31 DEGs associated with ABC transporters, six subfamilies were represented: ABCG5 (11 DEGs), ABCG2 (3 DEGs), ABCC4 (1 DEG), ABCC1 (9 DEGs), ABCB1 (2 DEGs), and ABCA3 (4 DEGs) (Fig. 6C). Notably, the majority of these genes exhibited a downregulated expression pattern. Gene Ontology (GO) enrichment analysis further showed that, within the cellular component (CC) category, DEGs were significantly enriched in terms related to integral and intrinsic membrane components as well as membrane-associated regions. Within the molecular function (MF) category, significant enrichment was observed for terms associated with transporter activity and transmembrane transporter activity (Fig. 6D).

**Fig. 6.**
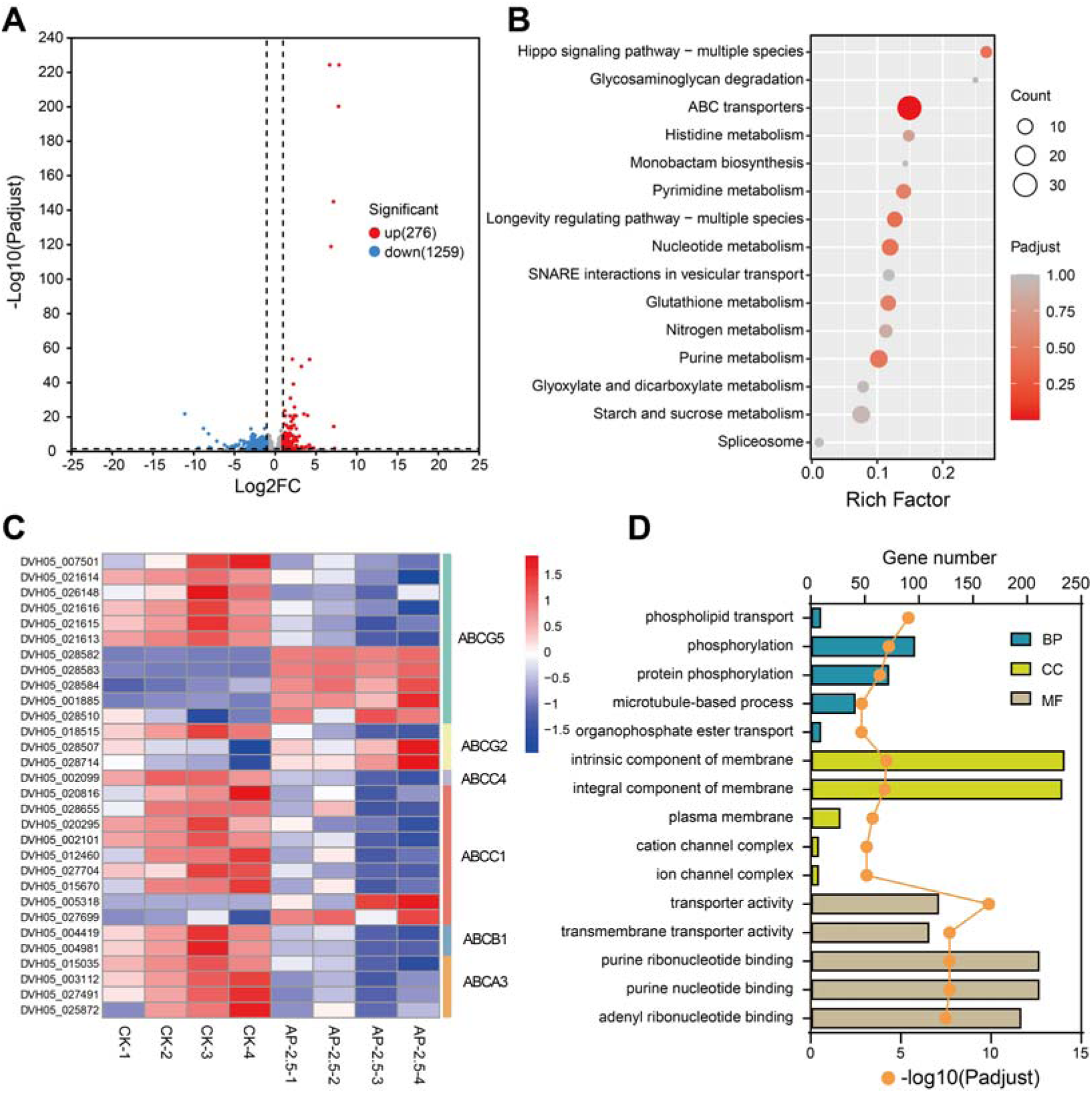
Transcriptomic analysis of *P. capsici* in response to apigenin treatment. (A) Volcano plots of DEGs in *P. capsici* under conditions of apigenin treatment. (B) KEGG enrichment analysis was performed on transcripts differentially expressed in *P. capsici* following apigenin treatment. (C) Heatmap analysis of the expression profiles of 31 genes encoding ABC transporters under control conditions and following apigenin treatment. (D) Go enrichment analysis was performed on transcripts differentially expressed in *P. capsici* following apigenin treatment.

## Discussion

We compared resistant and susceptible pepper cultivars and observed distinct response strategies to pathogen infection. Resistant plants maintained a relatively stable rhizosphere microbiome composition and metabolic profile (Fig. 2A, B), indicating a stronger capacity to sustain homeostasis under pathogen challenge. Similar patterns have been observed in other crops. For instance, a metabolomic study of tobacco challenged by *Ralstonia pseudosolanacearum* showed that the susceptible cultivar Yunyan87 exhibited significant changes in 88 known metabolites following infection, whereas the resistant cultivar K326 displayed almost no such changes. These findings are consistent with the Anna Karenina Principle (AKP) applied to plant microbiota: “All healthy microbiota are alike; each disease-associated microbiota is sick in its own way” [35].

We performed transcriptomic and metabolomic analyses of resistant pepper materials and found that the flavonoid pathway was strongly activated in the resistant cultivar (Fig. 2F, G; Fig. S2A, B). Flavonoids have long been recognized as secondary metabolites with antimicrobial activity [36,37]; for example, apigenin has been shown to inhibit the growth of several soybean pathogens [38]. In wheat, apigenin and its derivatives have also been demonstrated to play a pivotal role in resistance against *Fusarium* head blight [39]. More recently, increasing evidence has highlighted their ecological role in the rhizosphere: flavonoids such as apigenin and luteolin can recruit beneficial microbial taxa, including *Pseudomonas*, *Rhizobium*, and *Oxalobacteraceae*, thereby enhancing nutrient acquisition and promoting plant health [29]. Root-secreted metabolites not only provide nutrients for microbial communities but also act as selective cues that shape the assembly of beneficial consortia [40,41]. While numerous studies have demonstrated either the antimicrobial activity or the microbiome-modulating function of flavonoids individually, there are relatively few reports that simultaneously validate both roles within a single plant–pathogen–microbiome system. Our study in the pepper *P. capsici* system demonstrates this dual function of apigenin: it not only suppresses pathogen propagation by disrupting sporangia and inhibiting zoospore release, but also acts as a hub metabolite that recruits disease-suppressive microbes in the rhizosphere. These findings support the notion that flavonoids serve not only as “weapons” (antimicrobial agents) but also as a chemical mediator orchestrating interactions between the host plant and its surrounding microbiome.

PAL, C4H, 4CL, CHS, and CHI are key enzymes in the early stages of flavonoid biosynthesis and typically respond to pathogen invasion [42]. Specifically, C4H, CHS, CHI, and FNSI/FNSII direct metabolites in the phenylpropanoid pathway toward apigenin and luteolin, thereby positively regulating their accumulation. Our RNA-seq analysis revealed that in the resistant pepper line, C4H expression was upregulated at 4 days post-inoculation, while PAL and 4CL were upregulated at both 2- and 4-days post-inoculation; in contrast, these genes were mainly downregulated in the susceptible line (Fig. 3A). This suggests that resistant peppers are able to sustain the accumulation of precursors required for apigenin biosynthesis during infection, providing substrates for downstream flavonoid production. By contrast, the downregulation of these genes in susceptible peppers may limit precursor availability, potentially resulting in insufficient apigenin synthesis. CHS is the key entry enzyme in the apigenin biosynthetic pathway, catalyzing chalcone formation as the initial step in flavonoid synthesis. Reduced CHS expression or activity typically limits downstream flavonoid production, including apigenin [43]. Consistently, CHS transcript levels are positively correlated with apigenin content [44,45]. Notably, CHS expression was markedly reduced in the susceptible cultivar. (Fig. 3A), which indicates a constrained capacity for apigenin biosynthesis, potentially contributing to the weakened defense response observed in this cultivar. FNS genes catalyze the conversion of naringenin to apigenin, promoting apigenin accumulation [46–48]. In resistant peppers, although FNS expression was transiently downregulated at 2 days post-inoculation (Fig. 3A), it subsequently recovered and was upregulated, enabling increased apigenin synthesis to enhance defense responses. In contrast, susceptible peppers lack this regulatory mechanism; FNS expression remained downregulated, likely restricting apigenin biosynthesis and reducing their ability to resist pathogen infection.

The dual functionality of a single metabolite has also been documented in other crops. For example, in tobacco, root-secreted methyl-ferulate not only suppresses *Phytophthora nicotianae* but also enriches rhizosphere *Bacillus* populations, thereby enhancing resistance against black shank disease [15]. Similarly, in rice, 4-hydroxycinnamic acid (4-HCA), a lignin precursor, has been shown to selectively recruit beneficial phyllosphere *Pseudomonas* while simultaneously inhibiting pathogenic *Xanthomonas* populations. This dual role effectively curtails the occurrence of bacterial blight and maintains a “healthy state” of the rice phyllosphere [49]. Such evidence underscores the significance of multifunctional metabolites in plant defense, highlighting their potential to both directly suppress pathogens and modulate the plant-associated microbiome. These findings suggest that exploiting such dual-action metabolites could represent a promising strategy for sustainable disease management in crops.

Plants influence microbes through metabolic secretions, and microbes reciprocally strengthen plant defense, forming a metabolite–microbiome feedback loop. In our study, apigenin exemplifies such a selective metabolite, promoting the recruitment of disease-suppressive microbes and strengthening a protective rhizosphere community (Fig. S3). Previous studies have reported similar mechanisms with phenolic acids such as ferulic acid or Vanillic acid, which alter microbial community composition and suppress soil-borne pathogens [50–52]. Notably, our results indicate resistant pepper cultivars were enriched with *Streptomyces* species (Fig. 4A). This observation is consistent with the growing evidence that Streptomyces recruitment represents a common strategy by which plants harness beneficial microbes to counteract pathogens. Similar findings have been reported in tomato, where two key root exudates enhanced resistance to *Ralstonia solanacearum* by recruiting *Streptomyces* [53]. In rice, *Streptomyces* species have also been shown to trigger defense responses against *Xanthomonas oryzae* pv. *oryzae,* thereby enhancing resistance to bacterial blight [54]. Interestingly, in the context of potato common scab, nonpathogenic *Streptomyces* strains were identified as important antagonists that suppress disease development.

RNA-seq analysis revealed that apigenin treatment induced 1,535 differentially expressed genes (DEGs) in *P. capsici* (Fig. 6A), with most being downregulated. Notably, many DEGs were enriched in pathways related to membrane structure (Fig. 6D). Consistently, membrane integrity assays showed increased green fluorescence in apigenin-treated hyphae (Fig. 5G) and enhanced permeability (Fig. 5H), providing direct evidence of apigenin-induced membrane damage. Notably, DEGs were also significantly enriched in the ABC transporter pathway. ABC (ATP-binding cassette) transporters represent one of the largest families of transmembrane proteins [55] and are widely involved in the secretion of secondary metabolites. In plants, ABC transporters mediate the export of defense-related compounds such as isoflavonoids, polyacetylenes, and phytoalexins to sites of pathogen invasion [56]. *Oomycetes*, due to their limited capacity to degrade plant defense compounds, often rely on ABC transporters to export such toxic molecules and occasionally to secrete their own virulence factors [57]. Thus, suppression of ABC transporter expression by apigenin may impair *P. capsici* detoxification and virulence, thereby reinforcing host defense. This mechanism parallels previous observations that citral suppresses *P. capsici* by downregulating ABC transporters and disrupting membrane integrity [58].

In conclusion, this work identifies apigenin as a multifunctional metabolite that directly impedes pathogen reproduction and indirectly promotes the establishment of beneficial rhizosphere microbiomes (Fig. 7). By integrating transcriptomic, metabolomic, and microbiome analyses, our study illustrates how plant-derived metabolites function as central nodes in the network linking host defense, microbial recruitment, and disease suppression. These findings not only advance our mechanistic understanding of metabolite-microbiome-pathogen interactions but also highlight new opportunities for harnessing natural compounds to achieve sustainable crop protection.

**Fig. 7.**
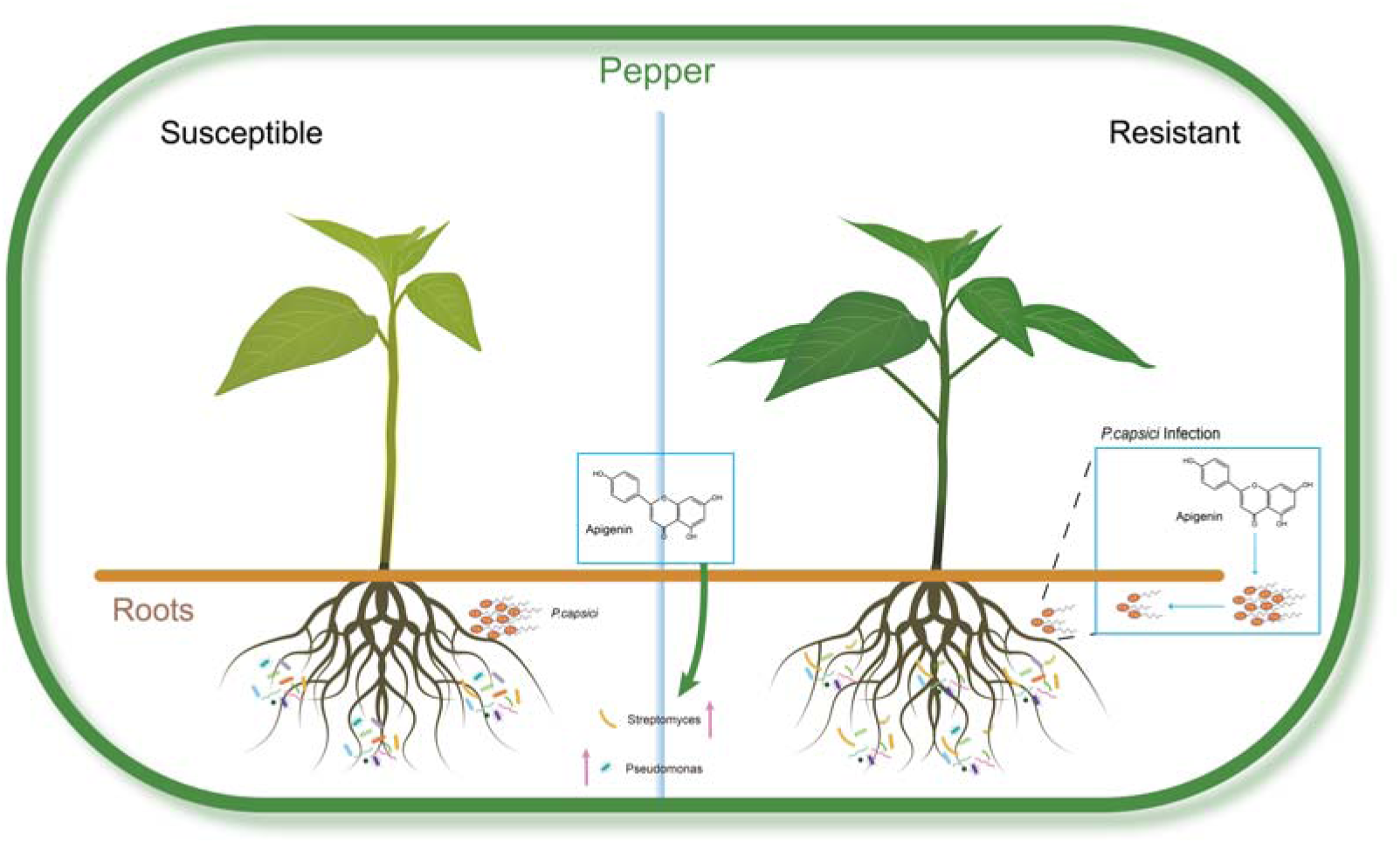
Schematic illustration of the proposed mechanism by which root-secreted apigenin enhances pepper resistance against *P. capsici*. The model depicts a dual mode of action: direct inhibition of *P. capsici* zoospore release and modulation of rhizosphere microbiome assembly.

## Materials and methods

### Plant materials and growth conditions

Two pepper (*Capsicum frutescens*) accessions were used in this study: the *P. capsici*–resistant accession CA53 and the susceptible accession CA476. Seeds were sown in a plant growth mix and maintained in a climate chamber at 28 °C under a 16 h light/8 h dark photoperiod. Fourteen days after sowing, seedlings were transplanted into individual 9-cm pots filled with the same mix soil. For co-cultivation treatment, one CA53 and one CA476 seedling were transplanted into the same pot at a spacing of 3–4 cm. Seedlings were used for inoculation when they reached the 6–8 true-leaf stage (approximately 4 weeks after sowing).

### Detached leaf inoculation (*in vitro*) and whole-plant inoculation (*in vivo*) with *P. capsici*

For *in vitro* assays, healthy, mature pepper leaves were surface sterilized (wiped with 75% ethanol followed by rinsing with sterile water), and 10 μL of zoospore suspension was placed at the middle of the leaf. The inoculated leaves were placed in Petri dishes lined with moist filter paper, sealed with plastic film to maintain humidity, and incubated at 25–28 °C. Disease symptoms were evaluated 3 days post-inoculation by photographing leaves and measuring lesion area through ImageJ.

For *in vivo* assays, seedlings at the 6–8 true-leaf stage were inoculated by drenching the root zone with 1 mL of the zoospore suspension (1 × 10L spores/mL). Disease severity was assessed 7 days post-inoculation based on a five-point scale [59]

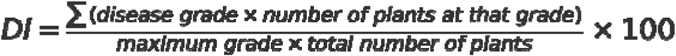

For the microbial transplantation group, pretreatment was performed 3 days prior to *P. capsici* inoculation. Specifically, rhizosphere soil of CA53 that had been inoculated with *P. capsici* was diluted at a soil-to-water ratio of 1:99. The diluted soil suspension was incubated in a shaker at 28°C with 200 rpm for 2 hours. Subsequently, 40 mL of the suspension was applied to each pot of CA476 plants as the microbial transplantation treatment. An additional application of the same prepared soil suspension (40 mL per pot) was repeated on the same day as *P. capsici* inoculation.

### Metabolite extraction and UPLC–MS/MS analysis

*Sample collection:* Metabolites were collected using a combination of in situ and hydroponic methods. Residual soil on roots was washed off with deionized water, and plants were placed in 40 mL of deionized water in centrifuge tubes. Roots were kept in the dark under their original growth conditions. After 24 hours, the solution was collected as root exudates, while roots were excised and used for endogenous metabolite analysis. All samples were stored at –80 °C prior to analysis.

*Roots metabolite extraction:* Plant samples were freeze-dried using a vacuum freeze dryer (Scientz-100F) and ground into fine powder with a mixer mill (MM 400, Retsch; 30 Hz, 1.5 min). A total of 50 mg powder was extracted with 1.2 mL of 70% methanol containing internal standards. Samples were vortexed every 30 min (30 s each time, six times in total), then centrifuged at 12,000 rpm for 3 min. The supernatant was filtered through a 0.22 μm microporous membrane and stored in sample vials for UPLC–MS/MS analysis.

*Root exudate extraction*: Samples stored at –80 °C were thawed on ice and vortexed for 10 s. A 9 mL aliquot was vacuum freeze-dried, and the dried material was resuspended in 300 μL of 70% methanol containing internal standards. The mixture was vortexed for 3 min and sonicated for 10 min, then centrifuged at 12,000 rpm for 10 min at 4 °C. The supernatant was filtered through a 0.22 μm microporous membrane and transferred into vials for UPLC–MS/MS analysis.

### Rhizosphere soil sampling and microbial community analysis

Rhizosphere soil samples were collected as follows. Plants were gently uprooted, and the loosely adhering soil was shaken off and discarded. The soil tightly adhering to the roots was then carefully brushed off using a sterile brush and collected. The collected rhizosphere soil was divided into two portions: one was stored at –80 °C for long-term preservation, and the other was kept at 4 °C for short-term storage.

Total microbial genomic DNA was extracted from soil samples using the E.Z.N.A.® soil DNA Kit (Omega Bio-tek, Norcross, GA, U.S.) according to manufacturer’s instructions. The quality and concentration of DNA were determined by 1.0% agarose gel electrophoresis and a NanoDrop2000 spectrophotometer (Thermo Scientific, United States) and kept at −80 L prior to further use. The V3-V4 region of the 16S rRNA gene was amplified by PCR using the primer pair 341F (5’-CCTAYGGGRBGCASCAG-3’) and 806R (5’- GGACTACNNGGGTATCTAAT -3’). Amplicon sequences were generated using a NovaSeq 6000 sequencer (Illumina) with paired-end 250 bp mode (PE250) at Novogene Bioinformatics Technology. Quality-filtered reads were loaded into the Qiime2 pipeline [60]. Raw forward and reverse sequencing reads were quality filtered with DADA2 [61]. Taxonomy was assigned to amplicon sequence variants (ASVs) using scikit-learn naïve Bayes taxonomy [62] classifier against the SILVA sequence database (v. 138) [63].

### Flavonoid treatment on pepper plants

Detached leaves of the susceptible cultivar CA476 were preconditioned under high-humidity conditions (relative humidity >90%) and uniformly sprayed with 1 mg/mL apigenin solution. Leaves treated with PEG400 served as negative controls. Subsequently, the leaves were inoculated with a zoospore suspension of *P. capsici* strain LT1534 (1 × 10L spores/mL). Inoculated leaves were placed in moist trays and incubated at 25 °C for 3 days. Lesion expansion areas were photographed and quantified using ImageJ software.

For whole-plant assays, pepper plants at the appropriate growth stage were treated twice with apigenin solutions (1 or 2.5 μg/mL) at 48 h intervals. Water and PEG400 treatments served as negative controls. Zoospore suspensions of LT1534 were prepared and adjusted to 1 × 10L spores/mL. Plants were inoculated using a rhizosphere inoculation method (1 mL suspension per plant). Three biological replicates were included. After inoculation, plants were maintained in a growth chamber at 25 °C with a 16 h light/8 h dark photoperiod and 60% ± 5% relative humidity. Disease resistance was assessed 7 days post-inoculation using the disease index (DI).

### Effects of apigenin on the cell membrane of *P. capsici*

Cell membrane integrity was examined using SYTOX™ Green Ready Flow™ (Thermo Fisher Scientific, Waltham, MA, USA) to stain *P. capsici* hyphae treated with apigenin, as previously described [58,64]. Membrane permeability was evaluated using a conductivity meter (Seven2Go Pro™, China) following apigenin treatment, as described previously [65,66]. Glycerol content was determined with a commercial glycerol assay kit (Nanjing Jiancheng Bioengineering Institute, Nanjing, China), following established protocols [50]. Each assay was performed with three biological replicates.

### Flavonoid treatment of *P. capsici* and transcriptomic analysis via RNA-seq

Mycelial plugs of LT1534 were transferred from solid medium to V8 liquid medium and incubated at 25 °C in the dark for 3 days. Mycelia were washed three times with sterile tap water (10 min each), and apigenin was added to a final concentration of 2.5 μg/mL. PEG400 treatment served as the negative control. Treated mycelia were incubated under dark conditions at 25 °C for 12 h to induce zoospore release. Samples were harvested for RNA-seq analysis, with four biological replicates per treatment.

### Statistical analysis

All statistical analyses were performed in R (v4.2.3, http://www.rproject.org/). Community alpha-diversity indexes based on 16S rRNA gene amplicon sequencing data were calculated with the ‘‘vegan’’ package (https://github.com/vegandevs/vegan). The beta-diversity was analyzed and visualized using principal coordinates analysis (PCoA) based on Bray-Curtis dissimilarity. Principal Component Analysis (PCA) was performed using the built - in prcomp function in R. Spearman rank correlation analysis of microbiome data was conducted using the corAndPvalue function implemented in the WGCNA package (v.1.73) [67]. All resulting networks were visualized using Gephi software (version 0.10.1) with the Fruchterman-Reingold layout algorithm to optimize node distribution and edge visualization, enhancing the clarity of network topological structures [68]. Prior to statistical analysis, all numeric variables in the dataset underwent log10 transformation to normalize their distribution, implemented using the dplyr package in R software (v4.3.0). One-way analysis of variance (ANOVA) was performed via the base R aov function. When significant group effects were detected, post-hoc comparisons were conducted using Tukey’s Honest Significant Difference (HSD) test with the TukeyHSD function. Graphical visualization was performed using the ggplot2 package (v3.4.4) in R software.

## Acknowledgements

We thank Dr. Qunqing Wang of Shandong Agricultural University for providing *P. capsici* isolate LT1534. We thank Dr. Taotao Wang and Mr. Mengzhuo Zheng help analyzed the transcriptome data. We thank Mass Spectrometry (Proteomics and Metabolomics) Platform of Peking University Institute of Advanced Agricultural Sciences for data determination. We thank Dr. Zhangsheng Zhu of South China Agricultural University and Dr. Hang He, Jianxin Bian of Peking University Institute of Advanced Agricultural Sciences for providing pepper materials.

## Authors’ contributions

Y.C. conceived and supervised the study. X.L. and W.S. designed the research. X.L., W.S., J.Z., Y.W., and P.R. conducted the experiments. X.L., W.S., and Y.W. analyzed the data. X.L., W.S., and Y.C. drafted the manuscript. All authors revised and approved the final version of the manuscript.

## Data availability

Transcriptome data of pepper roots, transcriptome data of *P. capsici* treated with apigenin, and soil microbiome data have been deposited in the Genome Sequence Archive of China National Center for Bioinformation (CNCB) (https://www.cncb.ac.cn/) under the BioProject PRJCA046279-CRA030058, CRA030168, CRA030165.

## Declarations

### Ethics approval and consent to participate

Not applicable.

### Consent for publication

Not applicable.

## Additional information

### Supplementary Information

Additional File 1. Fig. S1 Disease index of 16 different pepper cultivars.

Fig. S2 Comparative transcriptome analysis of resistant and susceptible pepper under Apigenin treatment.

Fig. S3 Accumulation of flavonoid metabolites in pepper roots under different treatments.

Fig. S4 Effects of different flavonoid compounds on *P. capsici*.

Table S1. Information of pepper species used in this study.

Additional File 2. Table S2. 1041 metabolites were identified in the samples of root exudates via.

Additional File 3. Supplementary video1. The effects of PEG400 on zoospore release in *P. capsici*.

Additional File 4. Supplementary video2. The effects of luteolin on zoospore release in *P. capsici*.

Additional File 5. Supplementary video3. The effects of apigenin on zoospore release in *P. capsici*.

Additional File 6. Supplementary video4. The effects of apigenin-7-glucoside on zoospore release in *P. capsici*.

Additional File 7. Supplementary video5. The effects of CK (H_2_O) on zoospore release in *P. capsici*.

Additional File 8. Supplementary video6. The effects of PEG400 on zoospore release in *P. capsici*.

Additional File 9. Supplementary video7. The effects of 1μg/mL of apigenin on zoospore release in *P. capsici*.

Additional File 10. Supplementary video8. The effects of 2.5μg/mL of apigenin on zoospore release in *P. capsici*.

## References

1. Qin C, Yu C, Shen Y, Fang X, Chen L, Min J, et al. Whole-genome sequencing of cultivated and wild peppers provides insights into capsicum domestication and specialization. Proc Natl Acad Sci. 2014; 111:5135–5140.

2. Venkatesh J, Lee S-Y, Back S, Kim T-G, Kim GW, Kim J-M, et al. Update on the genetic and molecular regulation of the biosynthetic pathways underlying pepper fruit color and pungency. Curr Plant Biol. 2023; 35–36:100303.

3. Sanogo S, Lamour K, Kousik CS, Lozada DN, Parada-Rojas CH, Quesada-Ocampo LM, et al. *Phytophthora capsici*, 100 Years Later: Research Mile Markers from 1922 to 2022. Phytopathology. 2023; 113:921–930.

4. Bulgarelli D, Garrido-Oter R, Münch PC, Weiman A, Dröge J, Pan Y, et al. Structure and function of the bacterial root microbiota in wild and domesticated barley. Cell Host Microbe. 2015; 17:392–403.

5. Hassani MA, Durán P, Hacquard S. Microbial interactions within the plant holobiont. Microbiome. 2018; 6:58.

6. Trivedi P, Leach JE, Tringe SG, Sa T, Singh BK. Plant–microbiome interactions: from community assembly to plant health. Nat Rev Microbiol. 2020; 18:607–621.

7. Xun W, Liu Y, Ma A, Yan H, Miao Y, Shao J, et al. Dissection of rhizosphere microbiome and exploiting strategies for sustainable agriculture. New Phytologist. 2024; 242:2401–2410.

8. Yin J, Zhang Z, Zhu C, Wang T, Wang R, Ruan L. Heritability of tomato rhizobacteria resistant to Ralstonia solanacearum. Microbiome. 2022; 10:227.

9. Ankati S, Podile AR. Metabolites in the root exudates of groundnut change during interaction with plant growth promoting rhizobacteria in a strain-specific manner. J Plant Physiol. 2019; 243:153057.

10. Upadhyay SK, Srivastava AK, Rajput VD, Chauhan PK, Bhojiya AA, Jain D, et al. Root exudates: mechanistic insight of plant growth promoting rhizobacteria for sustainable crop production. Front Microbiol. 2022; 13:916488.

11. O’Neal L, Vo L, Alexandre G. Specific root exudate compounds sensed by dedicated chemoreceptors shape azospirillum brasilense chemotaxis in the rhizosphere. Appl Environ Microbiol. 2020; 86:e01026–20.

12. Saleh D, Sharma M, Seguin P, Jabaji S. Organic acids and root exudates of brachypodium distachyon: effects on chemotaxis and biofilm formation of endophytic bacteria. Can J Microbiol. 2020; 66:562–575.

13. Balyan G, Pandey AK. Root exudates, the warrior of plant life: Revolution below the ground. South African Journal of Botany. 2024; 164:280–287.

14. Liu C, Geng H, Li W, Li Y, Lu Y, Xie K, et al. Innate root exudates contributed to contrasting coping strategies in response to *Ralstonia solanacearum* in resistant and susceptible tomato cultivars. J Agric Food Chem. 2023; 71:20092–20104.

15. Ma S, Chen Q, Zheng Y, Ren T, He R, Cheng L, et al. A tale for two roles: root-secreted methyl ferulate inhibits *P. nicotianae* and enriches the rhizosphere *bacillus* against black shank disease in tobacco. Microbiome. 2025; 13:33.

16. Rolfe SA, Griffiths J, Ton J. Crying out for help with root exudates: adaptive mechanisms by which stressed plants assemble health-promoting soil microbiomes. Curr Opin Microbiol. 2019; 49:73–82.

17. Wang Z, Song Y. Toward understanding the genetic bases underlying plant-mediated “cry for help” to the microbiota. iMeta. 2022; 1:e8.

18. De Coninck B, Timmermans P, Vos C, Cammue BPA, Kazan K. What lies beneath: belowground defense strategies in plants. Trends in Plant Science. 2015; 20:91–101.

19. Tsunoda T, van Dam NM. Root chemical traits and their roles in belowground biotic interactions. Pedobiologia. 2017; 65:58–67.

20. Zhang C, Feng C, Zheng Y, Wang J, Wang F. Root exudates metabolic profiling suggests distinct defense mechanisms between resistant and susceptible tobacco cultivars against black shank disease. Front Plant Sci. 2020; 11:559775.

21. Kasteel M, Ketelaar T, Govers F. Fatal attraction: How *Phytophthora* zoospores find their host. Seminars in Cell & Developmental Biology. 2023; 148–149:13–21.

22. Morris PF, Bone E, Tyler BM. Chemotropic and contact responses of *Phytophthora sojae* hyphae to soybean isoflavonoids and artificial substrates1. Plant Physiol. 1998; 117:1171–1178.

23. Tyler BM, Wu M, Wang J, Cheung W, Morris PF. Chemotactic preferences and ptrain *Variation* in the response of *Phytophthora sojae* zoospores to host isoflavones. Appl Environ Microbiol. 1996; 62:2811–2817.

24. Bag S, Mondal A, Majumder A, Mondal SK, Banik A. Flavonoid mediated selective cross-talk between plants and beneficial soil microbiome. Phytochem Rev. 2022; 21:1739.

25. Vandeputte OM, Kiendrebeogo M, Rasamiravaka T, Stévigny C, Duez P, Rajaonson S, et al. The flavanone naringenin reduces the production of quorum sensing-controlled virulence factors in *Pseudomonas aeruginosa* PAO1. Microbiology. 2011; 157:2120–2132.

26. Vandeputte OM, Kiendrebeogo M, Rajaonson S, Diallo B, Mol A, El Jaziri M, et al. Identification of catechin as one of the flavonoids from combretum albiflorum bark extract that reduces the production of quorum-sensing-controlled virulence factors in *Pseudomonas aeruginosa* PAO1. Appl Environ Microbiol. 2010; 76:243–253.

27. Salehi B, Venditti A, Sharifi-Rad M, Kręgiel D, Sharifi-Rad J, Durazzo A, et al. The therapeutic potential of apigenin. Int J Mol Sci. 2019; 20:1305.

28. Banerjee K, Banerjee S, Das S, Mandal M. Probing the potential of apigenin liposomes in enhancing bacterial membrane perturbation and integrity loss. J Colloid Interface Sci. 2015; 453:48–59.

29. Wu J, Liu S, Zhang H, Chen S, Si J, Liu L, et al. Flavones enrich rhizosphere *Pseudomonas* to enhance nitrogen utilization and secondary root growth in populus. Nat Commun. 2025; 16:1461.

30. Coyte KZ, Schluter J, Foster KR. The ecology of the microbiome: Networks, competition, and stability. Science. 2015; 350:663–666.

31. Herren CM, McMahon KD. Cohesion: a method for quantifying the connectivity of microbial communities. ISME J. 2017; 11:2426–2438.

32. Mahmoudi M, Almario J, Lutap K, Nieselt K, Kemen E. Microbial communities living inside plant leaves or on the leaf surface are differently shaped by environmental cues. ISME Commun. 2024; 4:ycae103.

33. Yang C-X, Chen S-J, Hong X-Y, Wang L-Z, Wu H-M, Tang Y-Y, et al. Plant exudates-driven microbiome recruitment and assembly facilitates plant health management. FEMS Microbiol Rev. 2025; 49:fuaf008.

34. Ying W, Wang Y, Wei H, Luo Y, Ma Q, Zhu H, et al. Structure and function of the arabidopsis ABC transporter ABCB19 in brassinosteroid export. Science. 2024; 383:eadj4591.

35. Zaneveld JR, McMinds R, Vega Thurber R. Stress and stability: applying the anna karenina principle to animal microbiomes. Nat Microbiol. 2017; 2:17121.

36. Sun M, Li L, Wang C, Wang L, Lu D, Shen D, et al. Naringenin confers defence against *Phytophthora nicotianae* through antimicrobial activity and induction of pathogen resistance in tobacco. Mol Plant Pathol. 2022; 23:1737–1750.

37. Vijayakumar K, Ganesan V, Kannan S. Antibacterial and antibiofilm efficacy of quercetin against *Pseudomonas aeruginosa* and *Methicillin resistant* staphylococcus aureus associated with ICU infections. Biofouling. 2025; 41:211–224.

38. Jiang YN, Haudenshield JS, Hartman GL. Response of soybean fungal and oomycete pathogens to apigenin and genistein. Mycology. 2012; 3:153–157.

39. Su P, Zhao L, Li W, Zhao J, Yan J, Ma X, et al. Integrated metabolo-transcriptomics and functional characterization reveals that the wheat auxin receptor TIR1 negatively regulates defense against *Fusarium graminearum*. J Integr Plant Biol. 2021; 63:340–352.

40. Wen T, Yuan J, He X, Lin Y, Huang Q, Shen Q. Enrichment of beneficial cucumber rhizosphere microbes mediated by organic acid secretion. Hortic Res. 2020; 7:154.

41. Tong Y, Zheng X, Hu Y, Wu J, Liu H, Deng Y, et al. Root exudate-mediated plant–microbiome interactions determine plant health during disease infection. Agric Ecosyst Environ. 2024; 370:109056.

42. Hu X, Luo Z, Xu C, Wu Z, Wu C, Ebid MHM, et al. A comprehensive analysis of transcriptomics and metabolomics revealed key pathways involved in saccharum spontaneum defense against sporisorium scitamineum. J Agric Food Chem. 2024; 72:4476–4492.

43. Alam W, Rocca C, Khan H, Hussain Y, Aschner M, De Bartolo A, et al. Current status and future perspectives on therapeutic potential of apigenin: focus on metabolic-syndrome-dependent organ dysfunction. Antioxid (basel Switz). 2021; 10:1643.

44. Eloy NB, Voorend W, Lan W, Saleme M de LS, Cesarino I, Vanholme R, et al. Silencing CHALCONE SYNTHASE in maize impedes the incorporation of tricin into lignin and increases lignin content. Plant Physiol. 2017; 173:998–1016.

45. Yan J, Yu L, Xu S, Gu W, Zhu W. Apigenin accumulation and expression analysis of apigenin biosynthesis relative genes in celery. Sci Hortic. 2014; 165:218–224.

46. Righini S, Rodriguez EJ, Berosich C, Grotewold E, Casati P, Falcone Ferreyra ML. Apigenin produced by maize flavone synthase I and II protects plants against UV-B-induced damage. Plant Cell Environ. 2019; 42:495–508.

47. Lee YJ, Kim JH, Kim BG, Lim Y, Ahn J-H. Characterization of flavone synthase I from rice. BMB Rep. 2008; 41:68–71.

48. Luo J, Luo C, Han M, Wang Q, Song Z, Zhang H, et al. A natural variation of flavone synthase II gene enhances flavone accumulation and confers drought adaptation in chrysanthemum. New Phytol. 2025; 247:1445–1459.

49. Su P, Kang H, Peng Q, Wicaksono WA, Berg G, Liu Z, et al. Microbiome homeostasis on rice leaves is regulated by a precursor molecule of lignin biosynthesis. Nat Commun. 2024; 15:23.

50. Mendes R, Kruijt M, de Bruijn I, Dekkers E, van der Voort M, Schneider JHM, et al. Deciphering the rhizosphere microbiome for disease-suppressive bacteria. Science. 2011; 332:1097–1100.

51. Berendsen RL, Pieterse CMJ, Bakker PAHM. The rhizosphere microbiome and plant health. Trends in Plant Science. 2012; 17:478–486.

52. Wang B, Xia Q, Li Y, Zhao J, Yang S, Wei F, et al. Root rot-infected sanqi ginseng rhizosphere harbors dynamically pathogenic microbiotas driven by the shift of phenolic acids. Plant Soil. 2021; 465:385–402.

53. Yang K, Fu R, Feng H, Jiang G, Finkel O, Sun T, et al. RIN enhances plant disease resistance via root exudate-mediated assembly of disease-suppressive rhizosphere microbiota. Mol Plant. 2023; 16:1379–1395.

54. Ebrahimi-Zarandi M, Bonjar GHS, Riseh RS, El-Shetehy M, Saadoun I, Barka EA. Exploring two streptomyces species to control rhizoctonia solani in tomato. Agronomy. 2021; 11:1384.

55. OuYang Q, Tao N, Jing G. Transcriptional profiling analysis of *Penicillium digitatum*, the causal agent of citrus green mold, unravels an inhibited ergosterol biosynthesis pathway in response to citral. BMC Genomics. 2016; 17:599.

56. Judelson HS, Ah-Fong AMV. Exchanges at the plant-oomycete interface that influence disease. Plant Physiol. 2019; 179:1198–1211.

57. Ah-Fong AMV, Shrivastava J, Judelson HS. Lifestyle, gene gain and loss, and transcriptional remodeling cause divergence in the transcriptomes of *Phytophthora infestans* and pythium ultimum during potato tuber colonization. BMC Genomics. 2017; 18:764.

58. Cui K, Wang Y, Wang M, Zhao T, Zhang F, He L, et al. Inhibitory activity and antioomycete mechanism of citral against *Phytophthora capsici*. Pestic Biochem Physiol. 2024; 204:106067.

59. Li S, Zhihui C. Allium sativum extract as a biopesticide affecting pepper blight. Int J Veg Sci. 2008; 15:13–23.

60. Bolyen E, Rideout JR, Dillon MR, Bokulich NA, Abnet CC, Al-Ghalith GA, et al. Reproducible, interactive, scalable and extensible microbiome data science using QIIME 2. Nat Biotechnol. 2019; 37:852–857.

61. Callahan BJ, McMurdie PJ, Rosen MJ, Han AW, Johnson AJA, Holmes SP. DADA2: high-resolution sample inference from illumina amplicon data. Nat Methods. 2016; 13:581–583.

62. Pedregosa F, Varoquaux G, Gramfort A, Michel V, Thirion B, Grisel O, et al. Scikit-learn: machine learning in python. J Mach, Learn, Res. 2011; 12:2825–2830.

63. Quast C, Pruesse E, Yilmaz P, Gerken J, Schweer T, Yarza P, et al. The SILVA ribosomal RNA gene database project: Improved data processing and web-based tools. Nucleic Acids Res. 2013; 41:D590–596.

64. Li J, Bai X, Zhu G, Liu S, Liu C, Wu M, et al. Sanxiapeptin is an ideal preservative with a dual effect of controlling green mold and inducing systemic defense in postharvest citrus. Food Chem. 2024; 453:139669.

65. Zhang R, Xu Q, Zhang Y, Zhu F. Baseline sensitivity and toxic actions of prochloraz to sclerotinia sclerotiorum. Plant Dis. 2018; 102:2149–2157.

66. Sun L, Na R, Jiang C, Cui K, He Y, Zhao T, et al. Bioactivity and control efficacy of benzovindiflupyr against athelia rolfsii in China. Plant Dis. 2023; 107:2359–2364.

67. Langfelder P, Horvath S. WGCNA: An R package for weighted correlation network analysis. BMC Bioinf. 2008; 9:559.

68. Bastian M, Heymann S, Jacomy M. Gephi: an open source software for exploring and manipulating networks. Proc Int AAAI Conf Web Soc Media. 2009; 3:361–362.

